# Parallel cryo electron tomography on *in situ* lamellae

**DOI:** 10.1101/2022.04.07.487557

**Authors:** Fabian Eisenstein, Haruaki Yanagisawa, Hiroka Kashihara, Masahide Kikkawa, Sachiko Tsukita, Radostin Danev

## Abstract

*In situ* cryo electron tomography of cryo focused ion beam milled samples emerged in recent years as a powerful technique for structural studies of macromolecular complexes in their native cellular environment. The lamella-shaped samples, however, have a limited area and are created with a necessary pretilt. This severely limits the possibilities for recording tomographic tilt series in a high-throughput manner. Here, we utilise a geometrical sample model and optical image shift to record tens of tilt series in parallel, thereby saving time and gaining sample areas conventionally used for tracking of specimen movement. The parallel cryo electron tomography (PACE-tomo) method achieves a throughput faster than 5 min per tilt series and allows the collection of sample areas that were previously unreachable, thus maximising the amount of data from each lamella. Performance testing with ribosomes *in vitro* and *in situ* on state-of-the-art and general-purpose microscopes demonstrated the high-throughput and high-quality of PACE-tomo.

## Introduction

In recent years, cryo electron tomography (cryoET) has been rapidly gaining popularity in the field of structural biology. Its capability to resolve molecular structures in their native cellular environment will only become more important as the field progresses. Many advances in sample preparation, data collection and data processing, which are essential for the determination of high-resolution structures, have been made (Hylton and Swulius, 2021). Data collection for cryoET is slow compared to the streamlined single particle analysis (SPA) cryo electron microscopy (cryoEM) workflows that have become routine for the structure determination of purified proteins and macromolecular complexes (Assaiya et al., 2021; Danev et al., 2021a). Unlike SPA cryoEM, where each image yields hundreds to thousands of identical particles, cryoET requires careful collection of several tens of images of each sample area at different tilt angles (tilt series) to extract only tens to hundreds of particles. Conventional tilt series acquisition for cryoET requires a tracking area to ensure that the region of interest does not leave the field of view when the stage is tilted. Depending on the precision of the sample holder and the size of the field of view, such specimen shifts can be detrimental to the data and must be compensated. The tracking area, usually a few micrometres along the tilt axis to ensure equivalent sample behaviour, is used to measure the specimen shifts and correct them by applying image shift before exposing the target area. On a flat sample, where the z height does not vary along the tilt axis, this approach is, although slow, very robust and widely used (Hagen et al., 2017; Schur et al., 2016; Tegunov et al., 2021). Recently, improved cryo microscope stages allowed for developments in high-throughput data collection under less challenging conditions, such as on flat and homogenous samples and larger pixel sizes (Chreifi et al., 2019; Eisenstein et al., 2019). Nevertheless, the general throughput in cryoET remains at least an order of magnitude lower than in SPA cryoEM.

SPA cryoEM data collection has benefited immensely from using increasingly larger beam image shifts to collect more images per stage position thereby minimising the rate-limiting stage movement, drift settling and autofocus steps (Cheng et al., 2018; Weis and Hagen, 2020; Wu et al., 2019). It has now become standard to collect upwards of 300 micrographs per hour (Danev et al., 2021b). This was enabled by compensation of beam shift-induced beam tilt in software (Zivanov et al., 2018) and in hardware via microscope calibration (Weis and Hagen, 2020). Making use of beam image shift acquisition in tomography is more complicated because distances between targets change throughout the tilt series. This requires precise modelling of sample movements as has previously been done for relatively flat samples (Bouvette et al., 2021; Zheng et al., 2004). However, *in situ* cellular samples elucidating the interactions of macromolecular complexes with each other and with the cellular architecture are becoming increasingly popular (Albert et al., 2020; Gupta et al., 2021; Mahamid et al., 2016; Schaffer et al., 2019; Weiss et al., 2022). Unfortunately, these samples are neither homogeneous nor flat due to their more complex sample preparation methods. The thinness of the sample is a major factor in obtaining high-resolution structures using cryoET. The highest resolution of a macromolecular complex *in situ* was obtained from a bacterium less than two hundred nanometres thick (Tegunov et al., 2021). Most eukaryotic cells have sizes in the order of micrometres to tens of micrometres. Cryo focused ion beam (cryoFIB) milling is currently the method of choice for sample thinning and preparation for *in situ* cryoET (Marko et al., 2007; Medeiros et al., 2018; Schaffer et al., 2017). Using cryoFIB milling, it is possible to obtain electron transparent lamellae that are less than three hundred nanometres thick. This process is becoming more automated and the preparation of tens of lamellae in a single day is now possible (Buckley et al., 2020; Klumpe et al., 2021; Tacke et al., 2021; Zachs et al., 2020).

Several experimental challenges arise when collecting cryoET data on lamellae. Most regions of a lamella contain biological sample and therefore, it is desirable to avoid using tracking areas which will not yield data. Additionally, lamellae are created with a pretilt dictated by the geometry of the cryoFIB-milling process. Thus, if a lamella is not oriented with the milling direction perpendicular to the tilt axis of the microscope, the z height of the sample varies along the tilt axis making tracking and compensation inaccurate (Schaffer et al., 2017). This makes it difficult to optimise throughput for robust data collection on lamellae.

Here, we developed a parallel cryo electron tomography (PACE-tomo) acquisition scheme based on a geometric model of the sample and beam image shift to collect tens of tomograms in parallel on a single lamella in less than 5 min per tilt series. The number and position of targets on a lamella can be arbitrarily selected and will influence the time per tilt series necessary for data collection. PACE-tomo is written in Python and runs within the free data acquisition software SerialEM (Schorb et al., 2019). To assess data quality, we reconstructed a sub-nanometre subtomogram average of eukaryotic ribosomes from a single lamella. Furthermore, we used PACE-tomo to collect 81 tilt series of an *in vitro* sample in 6 hours yielding a subtomogram average of a bacterial ribosome at a resolution of 3.1 Å.

## Results and Discussion

### Parallel acquisition of tilt series away from the tilt axis

For modern state-of-the-art sample holders, mechanical imperfections are minimal and sample movement can be predicted reasonably well based on geometrical considerations alone (Zheng et al., 2004). Employing beam image shift for the acquisition of tilt series in a parallel manner emphasises the importance of an accurate 3D model of the sample relative to the tilt axis. It has been shown previously, that this approach is feasible for purified samples on a holey carbon support film (Bouvette et al., 2021).

There are several factors that must be considered when collecting tilt series away from the tilt axis on a cryoFIB-milled lamella with a pretilt φ (Fig 1). By approximating the lamella as a plane and defining the tilt axis and the electron beam along x- and z-axes, respectively, the offset of the target area from the tilt axis and the pretilt of the lamella can be separated in y- and z-components, y_0_ and z_0_ (Fig. 1b). y_0_ is defined for every position during target selection on the untilted sample. The offset in z is subject to an additional error in eucentricity (zeuc), which applies to all targets equally, and 3D sample deformations (zerr) affecting each target individually (Fig. 1c). The determination of the resulting sum of errors z_0_* is not straightforward. Hence, after initialisation based solely on lamella pretilt, z_0_* is iteratively updated for every target position using measured specimen shifts throughout the tilt series. The number of data points used for the estimation of z_0_* can be adjusted to emphasise the compensation of local tilt angle-dependent behaviour of some sample holders (Zheng et al., 2004). In practice, using the last four recorded specimen shifts for each branch of the tilt series yielded good results.

**Figure 1:**
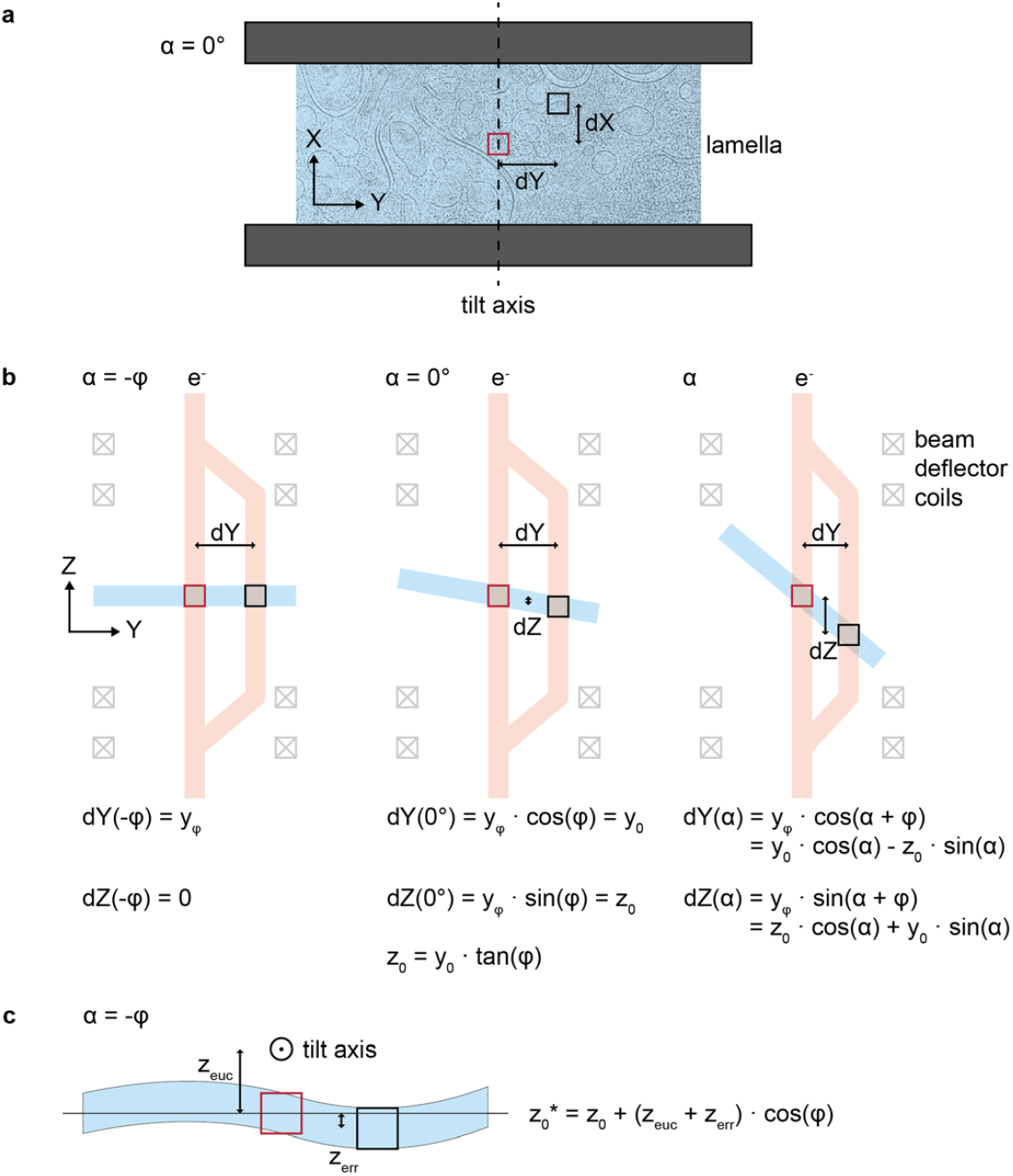
Lamella pretilt and sample geometry must be considered for accurate prediction of sample movement. **a**. Schematic of a cryoFIB-milled lamella in top view at a stage tilt angle α of 0°. The tracking target (red square) is on the tilt axis while an off-tilt axis target (black square) is displaced by dX and dY. **b**. Schematic of the acquisition geometry in side-view and equations to describe the position of the off-tilt axis target (black square) for different tilt angles. The tilt axis coincides with the tracking target (red square) (φ is the lamella pretilt angle, α is the stage tilt angle). **c**. Close-up of lamella showing 3D sample distortions and illustrating additional errors in eucentric height for all targets (zeuc) and per-target (z_err_).

### Speed and accuracy of PACE-tomo

We implemented PACE-tomo in a single Python script running within the scripting interface of SerialEM and tested it on the state-of-the-art cryo holder of a Krios G4 microscope (hereafter, “G4”) equipped with a Gatan K3 direct electron detector, as well as on room temperature and cryogenic side-entry holders in a JEM-F200 microscope (hereafter, “F200”) equipped with a Gatan K2 direct electron detector.

We used PACE-tomo routinely on the G4 for data collection on 41 lamellae with target numbers per lamella varying from 4 to 35. The tilt scheme was a dose-symmetric 120° tilt series with 3° increments, a starting tilt angle of 9° and a dataset-dependent exposure time. Fitting the collection times against the number of targets showed a typical collection time per tilt series of approximately 4.2 ± 0.1 min (Fig. 2a). Target locations can be arbitrarily set using a custom script that writes a target file with all necessary information and sets up the SerialEM navigator for batch acquisition of PACE-tomo areas. Figure 2b shows an example of 12 targets on a single lamella that were collected in 50 minutes. Targets were distributed over the whole lamella with beam-image shifts as far as 7 μm from the tilt axis (Fig. 2b). With the exception of the tracking tilt series (tilt series 1), the residual specimen shift errors along the y-axis (approximately perpendicular to the tilt axis) never exceeded ± 50 nm relative to the first tilt image in each tilt series (7.5 % of the field of view of a K3 detector at a pixel size of 1.64 Å) showing that despite the high throughput, field of view loss was minimal (Fig. 2c and d). The gain of field of view perpendicular to the tilt axis at higher tilt angles further compensated any displacements.

**Figure 2:**
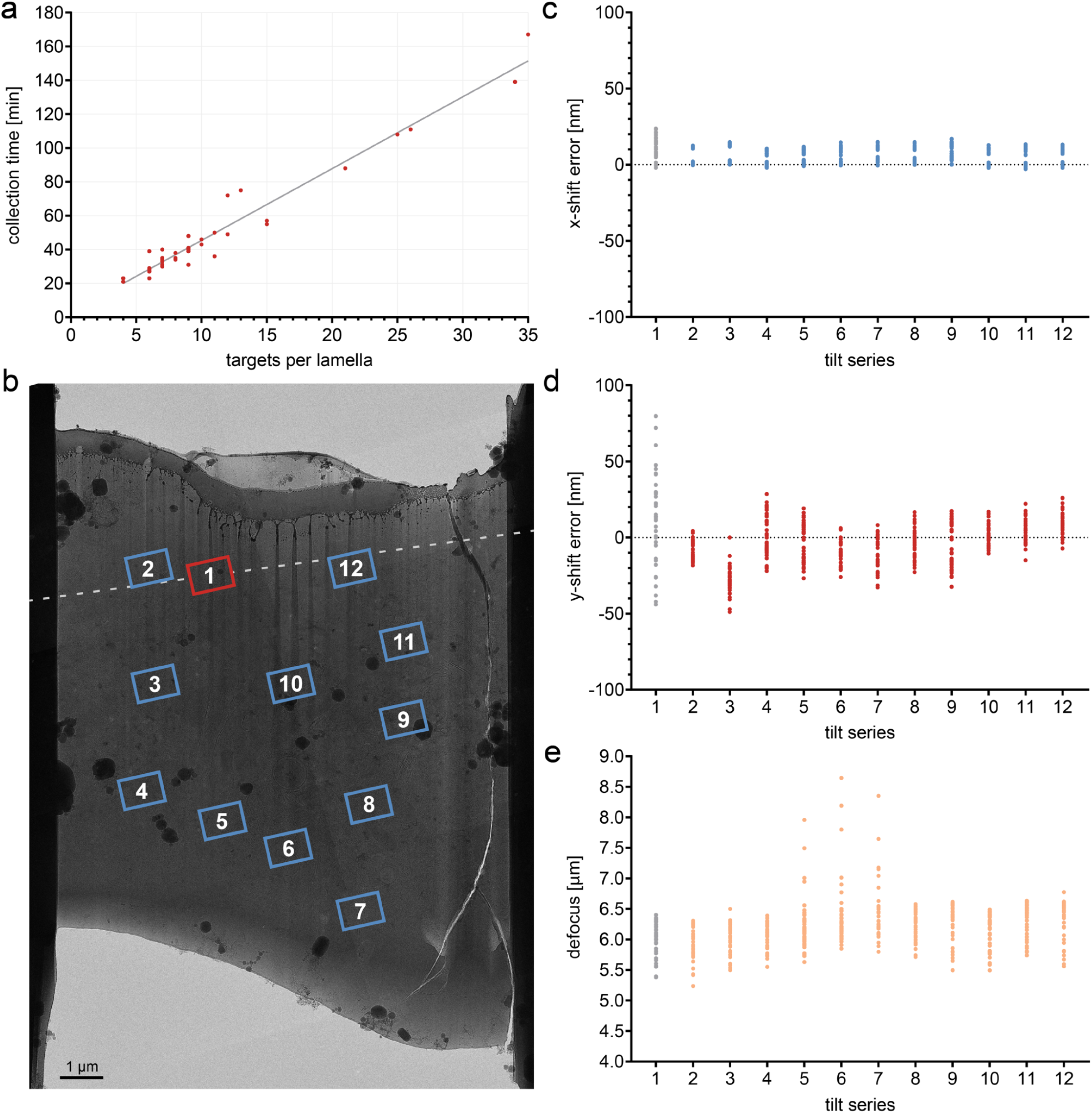
PACE-tomo keeps specimen shift errors within ± 50 nm. **a**. Collection time for 41 independent PACE-tomo runs with a varying number of targets per lamella. Linear regression shows a time per tilt series of 4.2 ± 0.1 min. **b**. An overview of a cryoFIB milled lamella with marked positions of tilt series. The targets are numbered according to their acquisition order. The red frame indicates the position of the tracking tilt series on the tilt axis (dashed line). Collection time was 50 min for 12 tilt series. **c and d**. Measured residual specimen shift errors by cross correlation-based alignment to the first image in each series in x- (**c**) and y-direction (**d**), parallel and perpendicular to the tilt axis, respectively. Tilt series 1 is the tracking tilt series with points coloured in grey. **e**. Defocus values for each tilt series estimated by CTF fitting. Four outliers in tilt series 7 caused by failed CTF fits due to large sample drift are not shown because they are outside of the plotted range.

The tracking tilt series is subject to the mechanical imperfections of the goniometer and hence suffers larger errors, which are corrected by an additional image shift applied before imaging the remaining targets. Furthermore, the tracking sample area is used for initial target alignment and autofocusing that lead to additional exposures. Nonetheless, specimen shift errors of the tracking tilt series stayed within ± 100 nm (15 % of the field of view of a K3 detector at a pixel size of 1.64 Å), keeping the loss of field of view minimal and the data usable (Fig. 2d). Shifts parallel to the tilt axis (x-shifts) were within ± 25 nm for all tilt series (Fig. 2c).

Residual errors showed no correlation with the distance from the tilt axis confirming the accuracy of the geometrical modelling. It is noteworthy, however, that the tilt axis offset from the optical axis must be determined precisely beforehand to ensure minimal errors in z-direction and hence, in defocus. The fine eucentricity routine implemented in SerialEM did not yield a sufficiently accurate estimate. A remaining tilt axis offset can be corrected by applying a linear defocus ramp vs tilt angle with an empirically determined slope (Supplementary Fig. 1). Using this approach, the defocus errors were kept within a range of approximately 1 µm (Fig. 2e). Outliers in the shown example are caused by failed CTF fitting due to factors like extensive drift, defocus gradient, and lower signal at higher tilt angles. Alternatively, we used a custom script to measure specimen shift and defocus change throughout a tilt series to solve for eucentric and tilt axis offset. The tilt axis offset remained constant over several imaging sessions.

Finally, to facilitate reliable tracking, features need to be discernible within the field of view of a given target. Failure of tracking is more likely at higher tilt angles due to lower contrast or target obstruction. The PACE-tomo routine worked well with a lamella thickness up to about 300 nm.

### PACE-tomo performance on a side-entry holder

To verify defocus compensation and investigate robustness of tracking against mechanical instability, we collected 13 dose-symmetric tilt series (± 60°) with an off-tilt axis image shift of up to 11.2 µm on a holey amorphous carbon layer on the F200 at room temperature (Fig. 3a). The collection time was about 82 min in total and about 6.3 min per tilt series.

**Figure 3:**
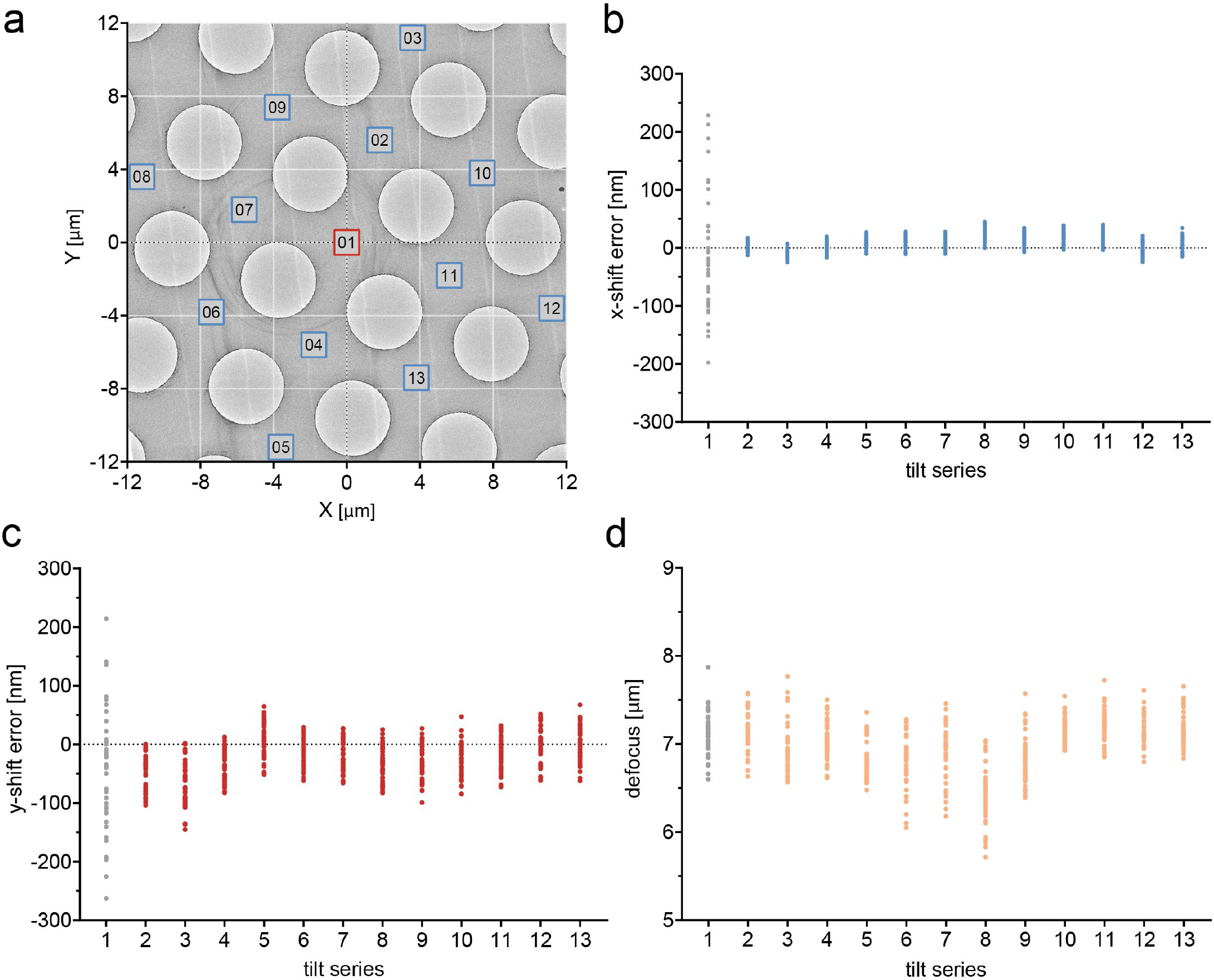
PACE-tomo performs well on a side-entry holder. **a**. An overview of a holey carbon sample area indicating the positions at which tilt series were collected. Targets are numbered according to acquisition order. Red frame indicates the position that was used for the tracking tilt series. The tilt axis is along Y=0. Collection time was 82 min for 13 tilt series. **b and c**. Measured residual specimen shift errors by cross correlation-based alignment to the first image (at 0°) in each tilt series in x- (**b**) and y-direction (**c**), parallel and perpendicular to the tilt axis, respectively. Tilt series 1 is the tracking tilt series and points are coloured grey. **d**. Defocus values for each tilt series estimated by CTF fitting.

Due to aberrant behaviour of the side entry holder depending on the positive and negative branch of the dose-symmetric tilt series (Supplementary Fig. 2), separate parameters were used for the tilt axis offset correction and additional backlash correction was performed for both branches. Taking these steps of extra precaution was essential for keeping the range of defoci throughout the tilt series contained. With exception of the tracking tilt series, remaining errors in specimen shifts were within ± 150 nm and the defoci were kept within a range of 1.5 µm.

We additionally tested PACE-tomo on the F200 using the Gatan 626 cryo-transfer holder, which only allowed for a tilt range of ± 30° at the chosen stage position. The collection time for 25 tilt series in a 5 by 5 equidistant pattern (Supplementary Fig. 3a) amounted to 118 min (∼4.7 min per tilt series). Given the restricted tilt range, specimen shift errors could be maintained within a range of ± 50 nm and ± 80 nm in x and y, respectively (Supplementary Fig. 3b and c). Defocus spread within each tilt series was about 0.5 µm (Supplementary Fig. 3d). The difference in defocus between tilt series shows that there was a considerable pretilt of the carbon support layer.

While the results are inferior to those of state-of-the-art microscope stages, in case of limited microscope access or for the purpose of screening a range of sample conditions, PACE-tomo makes high throughput data collection feasible on general purpose electron microscopes, like the JEM-F200.

### Subtomogram averaging of ribosomes *in vitro* and *in situ* using PACE-tomo

PACE-tomo significantly improves data collection throughput for structure determination of macromolecular complexes in their native environment. To illustrate its capabilities and to reaffirm that there is no loss in data quality, we collected 3 datasets and reconstructed subtomogram averages of ribosomes *in vitro* and *in situ* (Supplementary table 1). Datasets 1 and 2 were collected using purified 70S ribosomes *in vitro* on the G4 and the F200, respectively. Dataset 1 comprised 81 PACE-tomo tilt series in a 9 by 9 equidistant pattern (off-tilt axis image shifts up to 12 µm, Fig. 4a) required 6 hours on the G4 (∼4.5 min per tilt series). The number of target areas is limited by the amount of image shift that can be applied and the size of the grid square and grid bars. However, collecting data far from the tilt axis requires large defocus changes to compensate for changes in z-height at high tilt angles. Defocus values far from eucentric focus introduce additional optical aberrations that could be detrimental for the determination of high-resolution structures. Nevertheless, high-resolution information in tilt series is limited to the first tilt images due to accumulating radiation damage, hence, optical aberration effects at high tilt angles will be negligible in dose-symmetric tilt series.

**Figure 4:**
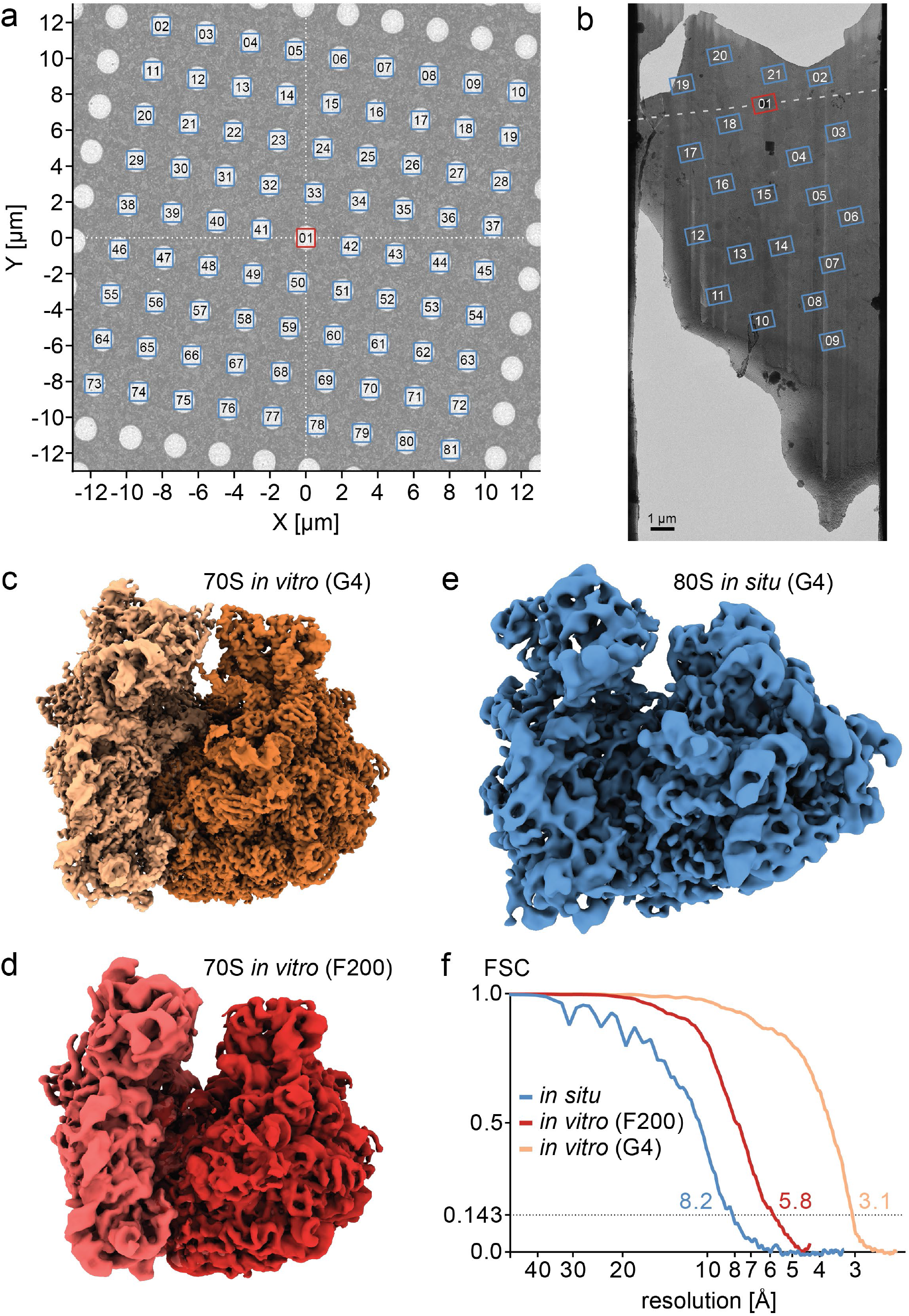
PACE-tomo datasets yield high resolution subtomogram averages. **a**. Acquisition pattern for dataset 1 – PACE-tomo with 81 targets in a regular 9 by 9 pattern. Targets are numbered according to acquisition order. The tracking tilt series (red frame) is at the origin and all other targets are acquired by relative image shift along the tilt axis (X-axis) and perpendicular to the tilt axis (Y-axis). Collection time was 6 hours. **b**. An overview of a cryoFIB-milled lamella indicating the positions at which tilt series for dataset 3 were collected. The red frame indicates the position of the tracking series on the tilt axis (dashed line) and targets are numbered according to acquisition order. Collection time was 88 min for 21 tilt series. **c**.3.1 Å subtomogram average of extracted 70S ribosomes from dataset 1 collected on the G4. Shown maps were filtered according to local resolution. Darker and lighter shades indicate large 50S subunit and small 30S subunit, respectively. **d**. 5.8 Å subtomogram average of extracted 70S ribosomes from dataset 2 collected on the F200 with a side-entry cryo-holder. The large 50S subunit (darker shade) and the small 30S subunit (lighter shade) were aligned separately. Shown maps were filtered according to local resolution. **e**.8.2 Å subtomogram average of *in situ* 80S ribosomes from dataset 3 collected on the G4. The shown map was filtered according to local resolution. **f**. Fourier Shell Correlation (FSC) curves of subtomogram averages. Resolution values were taken at a cut-off of 0.143.

Dataset 2 consisted of 25 tilt series from the F200 collected as described above with a tilt range of ± 30° (Supplementary Fig. 3). In contrast to the other datasets, dataset 3 collected on the G4 used *in situ* 80S ribosomes in their native cellular environment including ribosomes in the cytoplasm as well as attached to membranes (Supplementary Fig. 4a, Supplementary Movie 1). We collected 21 tilt series in 88 min (∼4.2 min per tilt series) on a single lamella with image shifts as far as 10 µm off the tilt axis (Fig. 4b).

Using these datasets, we determined subtomogram averages of 70S ribosomes *in vitro* at a resolution of 3.1 Å (dataset 1: 14,729 particles; Fig. 4c,f) and 5.8 Å (dataset 2: 10,540 particles; Fig. 4d,f). The subtomogram average of 80S ribosomes *in situ* yielded a resolution of 8.2 Å from 5,262 particles (Fig. 4e,f). Previously, subtomogram averaging of 80S ribosomes from cryoFIB-milled lamellae yielded resolutions in the range of 10-15 Å using considerably longer collection times (Pfeffer et al., 2017; Schaffer et al., 2019). Higher resolutions *in situ* have been achieved only for highly repetitive samples from tens of lamellae (Wang et al., 2022). Our results show that large beam image shifts and the higher collection speed of PACE-tomo do not compromise data quality, and the resulting tomograms are usable for studies of cellular architecture as well as subtomogram averaging. In fact, sample conditions in our example of 80S ribosomes *in situ* were less than ideal due to a partially broken lamella prone to beam-induced sample drift (Fig. 4b), as well as a lamella thickness of over 200 nm in some areas. It is noteworthy that PACE-tomo allows for the collection of sample areas that were previously out of reach due to a lack of space for focusing and tracking. This is often caused by partial loss of a lamella, target areas close to each other or close to the lamella edge, and previously resulted in an inability to acquire potentially valuable tomograms (Supplementary Fig. 4b).

## Conclusion

PACE-tomo allows simple and fast parallel collection of tomographic tilt series unconstrained by the tilt axis position. It works equally well on flat samples, allowing for fast collection of high-resolution structural data for purified proteins, and on complex geometries of cryoFIB-milled lamellae to study macromolecular complexes *in situ*. A possible limitation for the number of parallel acquisition targets is the amount of beam image shift that can be applied as well as radiation damage due to exposure overlap of neighbouring beam spots at high tilt angles. Furthermore, for the successful tracking of the target, sufficient signal for cross-correlation alignment must be present. This may affect excessively thick lamellae (>300 nm). Unlike recently published montage tomography workflows (Peck et al., 2021; Yang et al., 2022), PACE-tomo prioritises radiation damage avoidance, data quality, and throughput.

Together with the automation of lamella cryoFIB milling (Buckley et al., 2020; Tacke et al., 2021; Zachs et al., 2020), automated tilt series alignment and tomogram reconstruction (areTomo (Zheng et al., 2022), emClarity (Ni et al., 2022), dynamo (Burt et al., 2021)), PACE-tomo could play an important role in streamlining *in situ* cryoET workflows and determining high-resolution structures of macromolecular complexes in their native cellular environment.

## Acknowledgements

We are grateful to Prof. Yoshinori Fujiyoshi and Hiroshi Suzuki for stimulating discussions and valuable feedback. We thank Kazuhiko Nakamura and Yoichi Sakamaki for the management and support of the Graduate School of Medicine cryo-EM facility. We are grateful for support for cryoFIB milling and instrument maintenance by Yoshiyuki Fukuda. This research was partially supported by Platform Project for Supporting Drug Discovery and Life Science Research (Basis for Supporting Innovative Drug Discovery and Life Science Research (BINDS)) from AMED under Grant Number JP21am0101115j0005. F.E. is an International Research Fellow of the Japan Society for the Promotion of Science (JSPS, #P20764). H.Y. and M.K. were supported by Grant-in-Aid for Transformative Research Areas A (JSPS, 21H05248). S.T. received funding from Grant-in-Aid for Specially Promoted Research (JP19H05468). R.D. was supported by Takeda Science Foundation 2019 Medical Research Grant and Japan Science and Technology Agency PRESTO (18069571). This paper was typeset with the bioRxiv word template by @Chrelli: www.github.com/chrelli/bioRxiv-word-template

## Author contributions

F.E. developed PACE-tomo and conducted all experiments. H.Y. purified ribosomes and processed dataset 1. H.K. prepared the eukaryotic cell cultures. F.E., H.Y., H.K., M.K., S.T. and R.D. designed the experiments. F.E. wrote the manuscript with comments from all authors. Correspondence to

## Materials and Methods

### Cell culture and plunge freezing

Mouse mammary gland epithelial EpH4 cells were maintained in high-glucose Dulbecco’s modified Eagle’s medium (DMEM; Nissui Pharmaceuticals) supplemented with 10 % fetal bovine serum (FBS; Sigma) and 2 mM glutamine (Nacalai Tesque) at 37°C and 5 % CO2.

Electron microscopy grids were prepared as follows. Titanium grids (100 mesh) and gold grids (100 or 150 mesh) were placed on cover glass slides, coated with formvar-carbon and sterilised under ultraviolet light. EpH4 cells were carefully plated on the prepared grids in a 6-well plate at a density of 0.89×105 cells/cm^2^ to ensure that cells attached to only one side of the grids. The medium was changed daily for 7 to 9 days until EpH4 cell sheets had matured.

Sample grids were taken from cell culture wells and mounted in a Vitrobot Mark IV (Thermo Fisher Scientific). The sample chamber was kept at 37°C and 80 % humidity and grids were blotted for 10 s before being plunged into a liquid ethane-propane mixture. Grids were stored under liquid nitrogen until use.

### CryoFIB milling

Grids were clipped in modified autogrids with a cut-out to allow for shallower milling angles (Rigort et al., 2012). CryoFIB milling was done on an Aquilos 2 dual-beam instrument (Thermo Fisher Scientific). Samples were sputter-coated with Pt for 15 s at 30 mA and coated with a layer of organometallic Pt for 10 s using the gas injection system of the instrument. Targets were identified using the ion beam and initial 12 μm by 15 μm trenches above and below the target area were milled manually at high angles. Subsequently, MAPS and autoTEM (Thermo Fisher Scientific) were used to set up automated rough milling of the lamellae down to a target thickness of 250 nm reducing the ion beam current stepwise from 0.5 nA to 0.1 nA. Final polishing of the lamellae down to a target thickness of below 200 nm was performed manually with an ion beam current of 50 or 30 pA. Polishing times were kept below 1 h for all lamellae on a grid to reduce ice contamination build-up inside the cryoFIB chamber, which allowed the preparation of 5-10 lamellae per grid. Samples were removed from the holder and stored under liquid nitrogen.

### Ribosome purification and plunge freezing

70S ribosomes were prepared as described in (Ederth et al., 2009) with little modification. In brief, E. coli DH5alpha was used as host strain and the gene encoding the ribosomal protein L12 was tagged with a hexa-histidine affinity tag sequence at the C-terminus. HEPES-based Lysis buffer (30 mM HEPES-KOH pH 7.5, 10 mM MgCl2, 30 mM NH4Cl, 150 mM KCl, 1x EDTA-free protease inhibitor cocktail (Nacalai tesque)) was used for the purification. The purified His-tagged ribosomes were concentrated by centrifugation, resuspended in the lysis buffer, flash-frozen in LN2 and stored at -80°C before plunge freezing. Quantifoil Cu 200 R1.2/1.3 and UltrAuFoil Au 300 R1.2/1.3 grids were prepared by washing with acetone and glow discharging. Grids were frozen using a Vitrobot Mark IV (Thermo Fisher Scientific). The sample chamber was kept at 4°C and 100 % humidity and grids were blotted for 10 s before being plunged into a liquid ethane-propane mixture. Grids were stored under liquid nitrogen until use.

### PACE-tomo

Tilt series were recorded on a Krios G4 transmission electron microscope (Thermo Fisher Scientific) operated at 300 kV and equipped with a BioQuantum energy filter and K3 direct electron detector (Gatan). In case of cryoFIB-milled samples, grids were loaded with care to orient the lamellae perpendicular to the tilt axis. A custom Python script was used inside SerialEM 3.9 beta or 4.0 beta (Mastronarde, 2005; Schorb et al., 2019) to pick targets and set up the navigator. Tilt axis offset from the optical axis was determined experimentally and compensated by SerialEM internally. The sample was brought to eucentric height using the internal routine in SerialEM and the PACE-tomo script was run to collect selected targets in parallel. A simplified schematic illustrating the PACE-tomo method is shown in Supplementary Fig. 5. A dose-symmetrical tilt scheme covering 120° with 3° increments centred on a start tilt angle compensating for the lamellae pretilt (usually -9° depending on the orientation of the grid in the microscope), where necessary, was applied. More collection details for all datasets are shown in Supplementary Table 1.

PACE-tomo using a side-entry holder was performed on a JEM-F200 transmission electron microscope (JEOL) equipped with a 626 single tilt cryo-transfer holder (Gatan) and a K2 direct electron detector (Gatan). Tilt axis offsets were determined experimentally for both branches of the dose-symmetric tilt series to compensate for mechanical stage properties. A pixel size of 4.45 Å and a target defocus of -6 μm was used for testing tilt series on amorphous carbon, and 2.16 Å/pixel and a target defocus of -4 μm for dataset 2 of purified 70S ribosomes.

## Data processing and subtomogram averaging

Tilt series frames were aligned using *alignframes* in IMOD (Kremer et al., 1996; Mastronarde, 2008). Specimen shifts were determined by cross-correlation using *tiltxcorr* in IMOD and are given in image coordinates which are approximately parallel (x) and perpendicular (y) to the tilt axis.

Initial defocus estimation per tilt image was done by CTFFIND4 (Rohou and Grigorieff, 2015) or gctf (Zhang, 2016). Fine alignment of tilts was done by patch tracking and tomograms for particle picking were reconstructed by IMOD.

The subtomogram average workflows for all datasets are summarised in Supplementary Fig. 6.

Dataset 1: 20,663 particles were picked from 8x binned tomograms by *e2spt_tempmatch*.*py* in EMAN2 (Chen et al., 2019). Particle coordinates were imported to RELION-4.0-beta (Kimanius et al., 2021). After an initial 3D auto-refine of 8x binned pseudo-subtomograms followed by two rounds of 3D classification without alignment and removal of duplicate particles, 14,729 particles from classes resembling a 70S ribosome remained. Subsequent 3D auto-refine runs were performed using 4x and 2x binned pseudo-subtomograms and a local sampling range. Tomo CTF refinement (optics group per tilt series) and frame alignment were done in two rounds with one round of 3D auto-refine in between. The overall resolution of the 70S ribosome was 3.1 Å.

Dataset 2: 53,982 particles were picked from 8x binned tomograms by a custom-trained model using crYOLO (Wagner et al., 2019). Particle coordinates were imported into dynamo (Castaño-Díez et al., 2012) and an initial alignment using 8x binned subtomograms was performed. Particles were cleaned by cross-correlation coefficient and the remaining 22,441 particles were imported in RELION-4.0-beta using *dynamo2m* (Burt et al., 2021). After an initial 3D auto-refine of 8x binned pseudo-subtomograms followed by 3D classification without alignment and removing duplicate particles, 10540 particles from classes resembling a 70S ribosome remained. Subsequent 3D auto-refine runs were performed using 4x, 2x and unbinned pseudo-subtomograms and a local sampling range. Tomo CTF refinement (optics group per tilt series) and frame alignment were done in two rounds with one round of 3D auto-refine in between. Finally, masks for the 50S and 30S subunits were prepared and a local 3D auto-refine was performed separately yielding masked resolutions of 5.5 Å and 6.5 Å, respectively. The overall resolution of the 70S ribosome using the 50S masked alignment was 5.8 Å.

Dataset 3: 9,447 particles were picked by a custom-trained model using crYOLO. Particle coordinates were imported to RELION-4.0-beta and initial 3D auto-refine was performed using 4x binned pseudo-subtomograms. After 3D classification, 5,262 particles from classes resembling an 80S ribosome were selected. Following a 3D auto-refine with 2x binned pseudo-subtomograms, Tomo CTF refinement (optics group per tilt series) and Tomo frame alignment was performed. This was repeated one more round and the final reconstruction showed a resolution of 8.2 Å. Local resolution distributions were determined using RELION-4.0-beta. Structures were visualised using UCSF ChimeraX (Goddard et al., 2018). Tomograms were denoised for visualisation using cryo-CARE (Buchholz et al., 2018). Membrane-bound and cytoplasmic ribosomes were mapped into tomogram using the Python script *relionsubtomo2ChimeraX*.*py* (https://github.com/builab/subtomo2Chimera) and annotated manually.

## Data availability

Subtomogram averages were deposited in the Electron Microscopy Data Bank (EMDB accession codes: EMD-33115, EMD-33116, EMD-33117, EMD-33118), and tilt series in the Electron Microscopy Public Image Archive (EMPIAR accession codes: EMPIAR-10985, EMPIAR-10986, EMPIAR-10987).

SerialEM Python scripts for target selection and PACE-tomo are available at:

The SerialEM Script Repository (https://serialemscripts.nexperion.net/) and on https://github.com/eisfabian/PACEtomo.

**Supplementary Figure 1:**
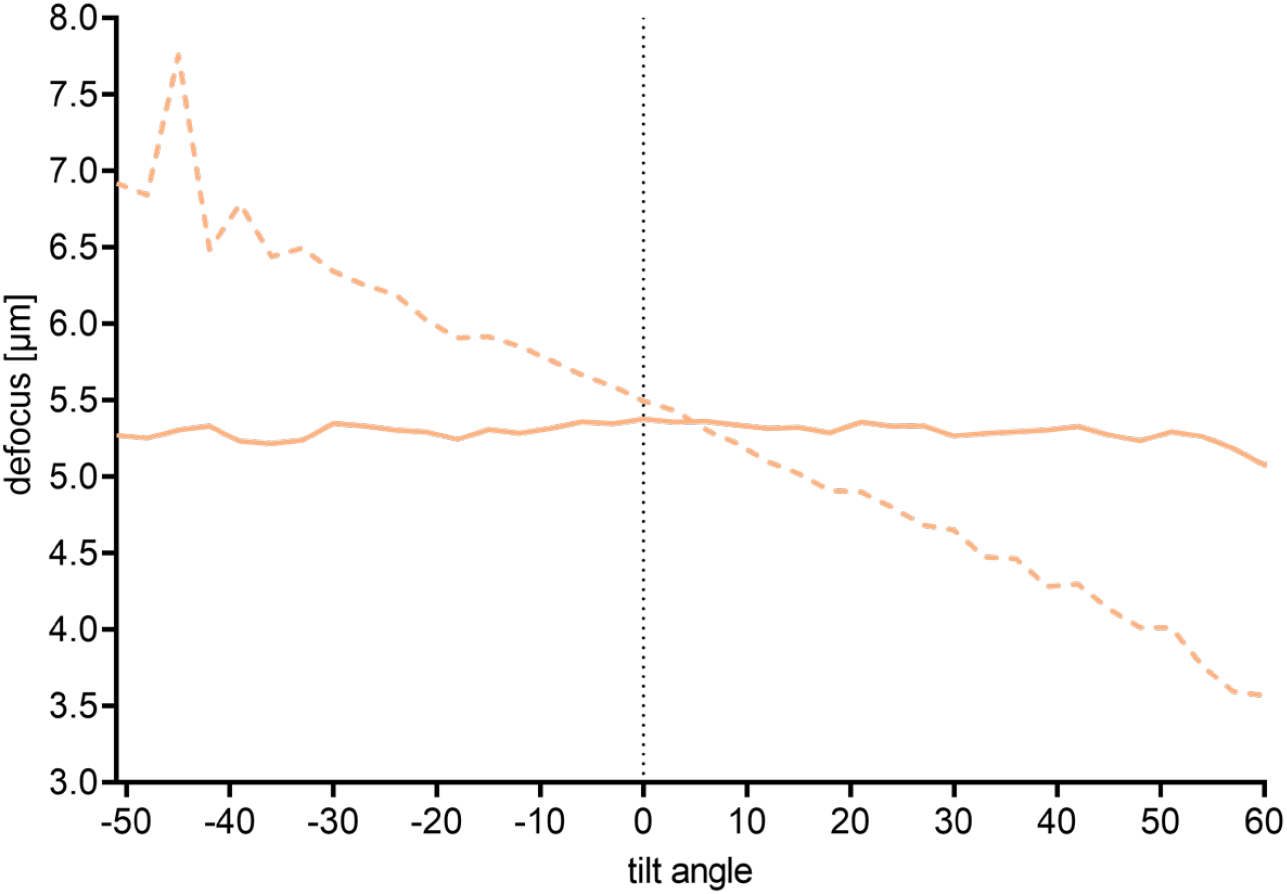
A linear defocus ramp can compensate for errors in tilt axis offset. Shown are defocus values estimated by CTF fitting for two tilt series collected on the same lamella. One tilt series was recorded without applying a defocus ramp (dashed line). The other tilt series (solid line) was recorded while applying a defocus ramp of 0.03 µm/degree.

**Supplementary Figure 2:**
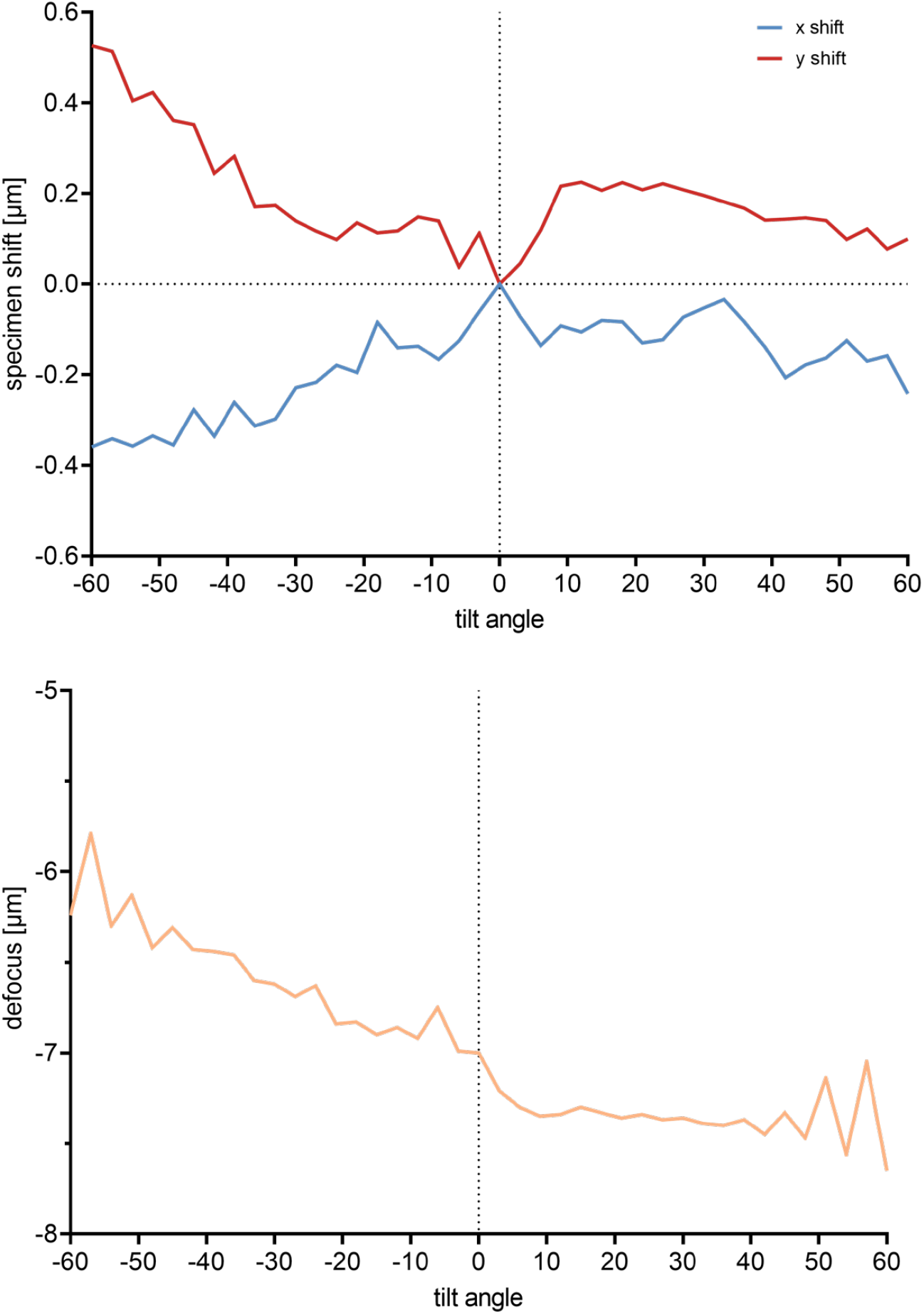
Side entry holder behaviour throughout dose-symmetric tilt series. Top: Specimen shifts in µm along x and y (parallel and perpendicular to the tilt axis respectively). The behaviour of the negative tilt angle branch differs from the positive tilt angle branch. Bottom: Defocus throughout the tilt series measured by beam tilt. While defocus remains relatively constant for the positive tilt angle branch, it shows a significant slope for the negative tilt angle branch.

**Supplementary Figure 3:**
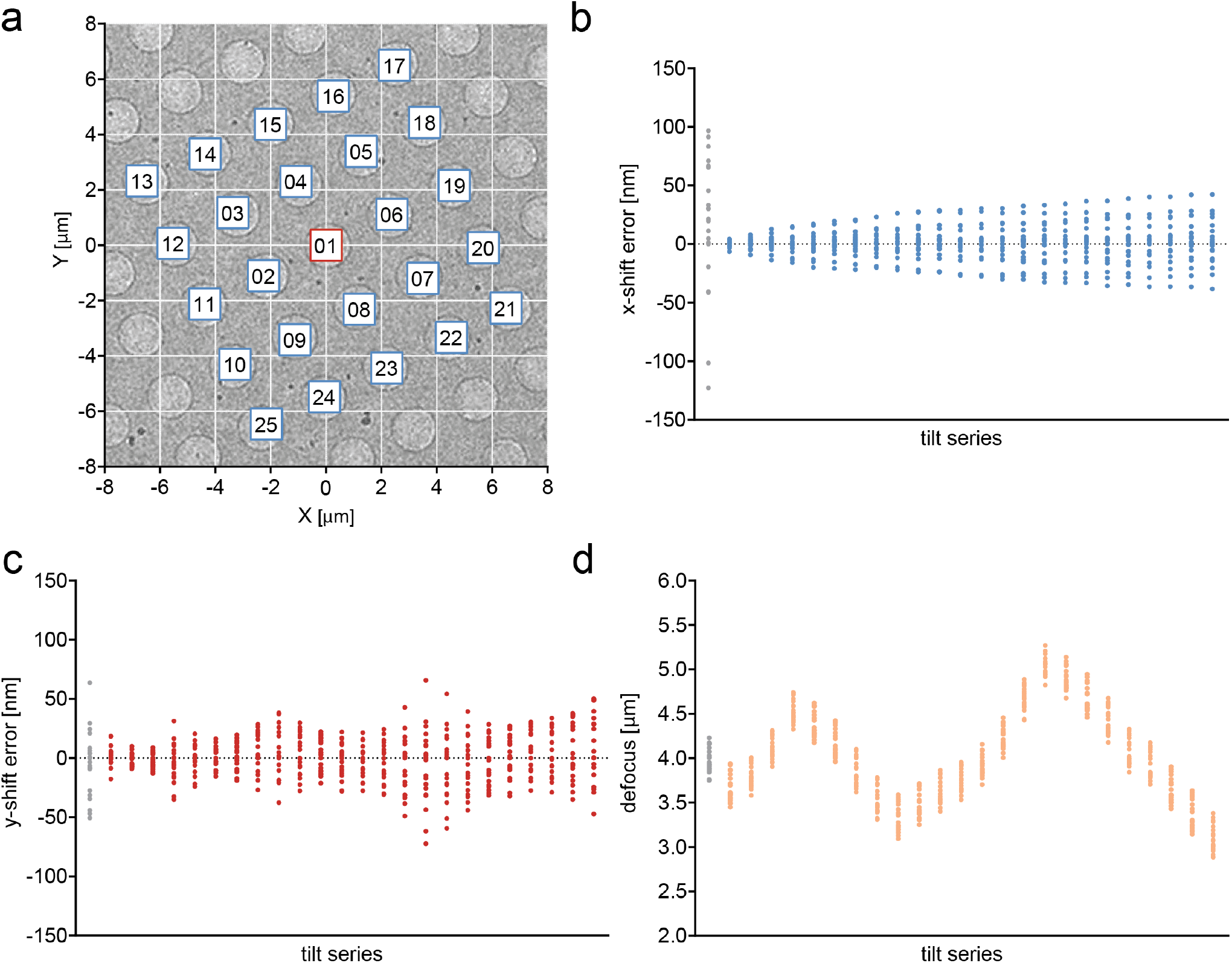
PACE-tomo tilt series collected using a cryo side-entry holder. **a**. An overview of a holey carbon sample area indicating the positions at which tilt series were collected. Targets are numbered according to acquisition order. Red frame indicates the position of the tracking tilt series. The tilt axis is along Y=0. Collection time was 118 min for 25 tilt series with an angular range of ± 30°. **b and c**. Measured specimen shift errors by cross correlation-based alignment to first tilt image at 0° in each tilt series in x- (**b**) and y-direction (**c**), parallel and perpendicular to the tilt axis, respectively. Tilt series 1 is the tracking tilt series and points are coloured grey. **d**. Defocus values for each tilt series estimated by CTF fitting.

**Supplementary Figure 4:**
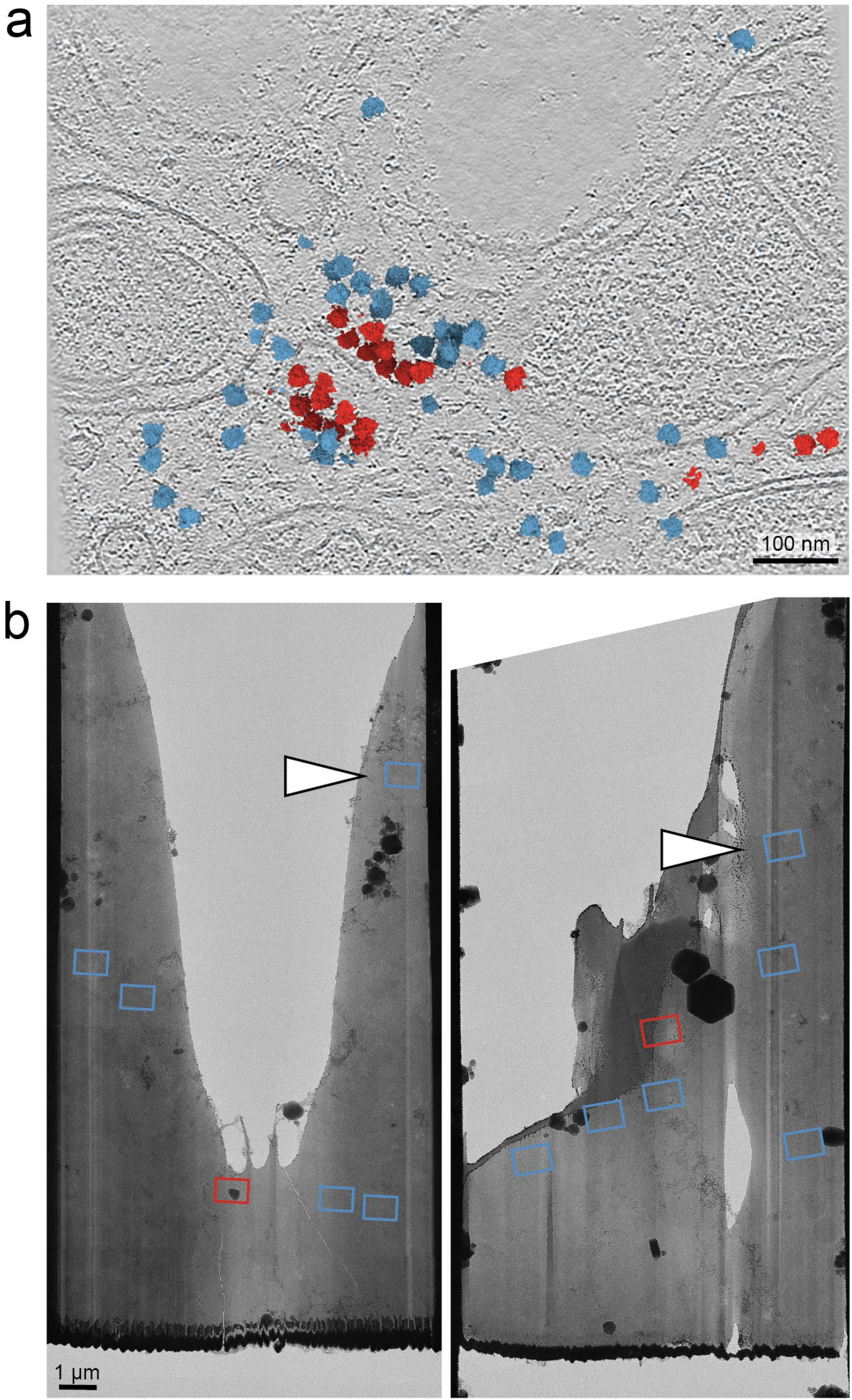
PACE-tomo facilitates data collection in challenging areas. **a**. Shown is a 0.66 nm thick slice through a cryo electron tomogram collected on the lamella shown in Fig. 4b. The subtomogram average was mapped into the tomogram using the refined coordinates and orientations. Membrane associated and cytoplasmic ribosomes are coloured red and blue, respectively. Scale bar: 100 nm. **b**. Overview of two cryoFIB-milled lamellae indicating the positions at which tilt series were collected. Red frame indicates the position that was used for the tracking tilt series. White arrowheads mark targets that would not have been accessible by conventional cryoET collection schemes. Collection times were 23 min for 6 tilt series (left) and 30 min for 7 tilt series (right).

**Supplementary Figure 5:**
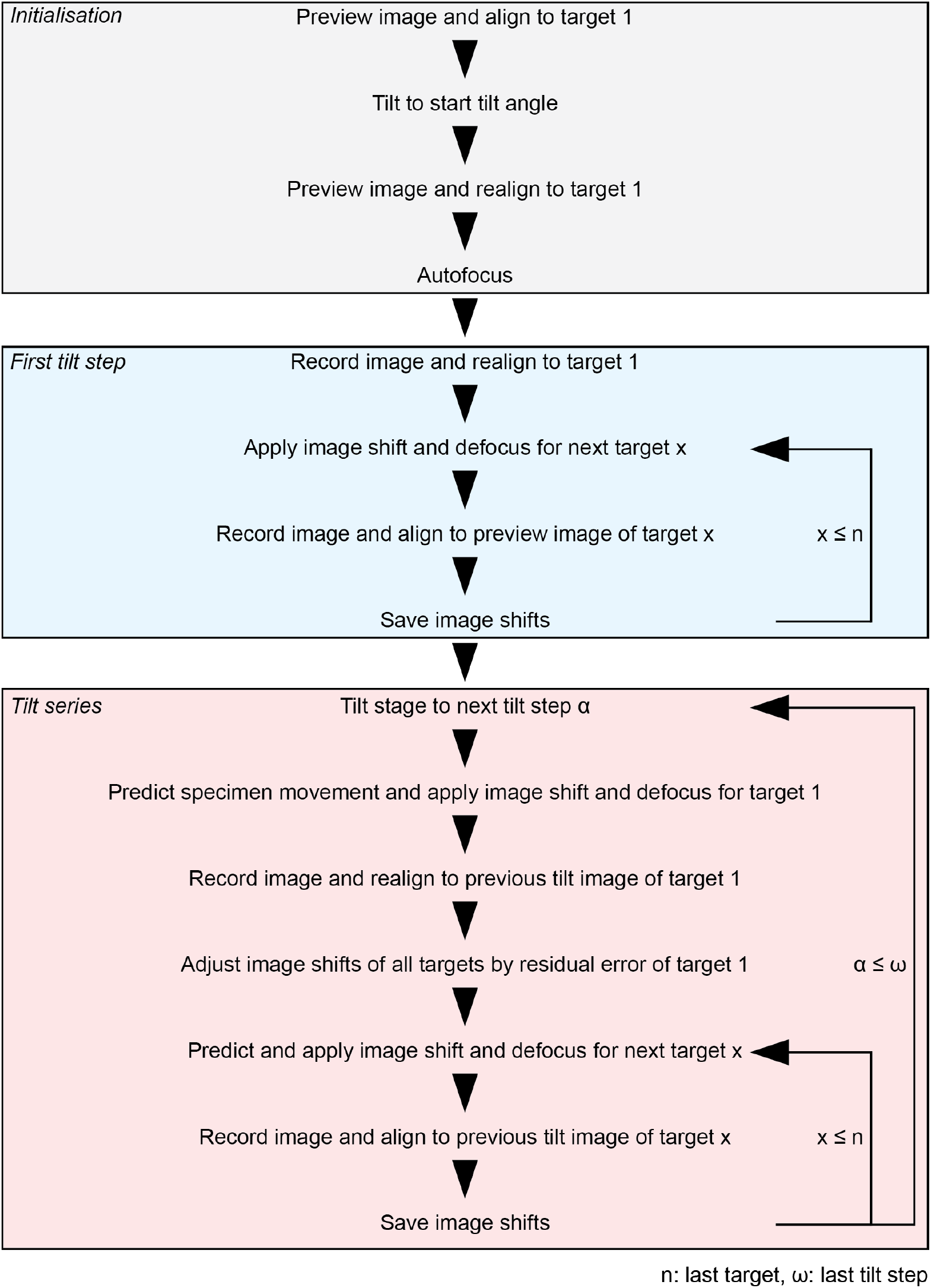
Simplified schematic of PACE-tomo. Shown are the essential steps of the PACE-tomo data acquisition scheme.

**Supplementary Figure 6:**
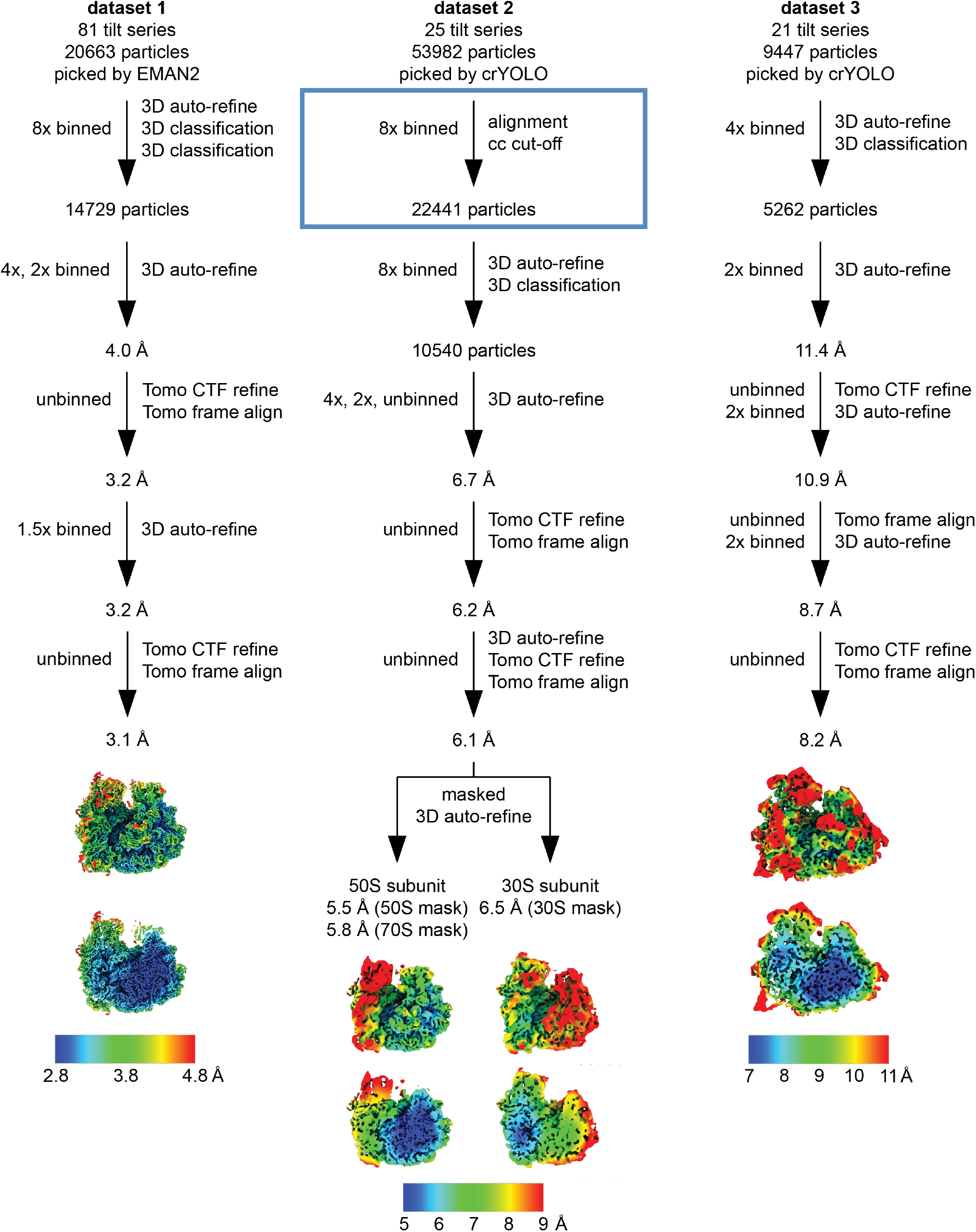
Processing workflows. Shown are the processing workflows for datasets 1-3 using RELION-4.0-beta. Blue box indicates steps done in dynamo. Final maps were filtered and coloured according to the local resolution distribution.

**Supplementary Table 1:** Collection parameters for ribosome datasets.

| Dataset |  | 1 | 2 | 3 |
| --- | --- | --- | --- | --- |
| Sample |  | 70S ribosomes<br>*in vitro* | 70S ribosomes<br>*in vitro* | 80S ribosomes<br>*in situ* |
| Microscope |  | Krios G4 | JEM-F200 | Krios G4 |
| Voltage (kV) |  | 300 | 200 | 300 |
| Camera |  | K3 | K2 | K3 |
| Energy filter slit width [eV] |  | 10 | None | 10 |
| Pixel size (Å) |  | 1.07 | 2.16 | 1.64 |
| Tilt range (°) |  | ± 60 | ± 30 | ± 60 |
| Tilt increment (°) |  | 3 | 3 | 3 |
| Total exposure (e<sup>-</sup>/Å<sup>2</sup>) |  | 177.3 | 113.4 | 144.8 |
| Movie frames per tilt image |  | 10 | 14 | 5 |
| Number of tilt series |  | 81 | 25 | 21 |
| Time per tilt series (min) |  | ~4.5 | ~4.7 | ~4.2 |
| Target defocus (µm) |  | -2.5 | -4.0 | -5.0 |
| Initial particles picked |  | 20,663 | 53,982 | 9,447 |
| Final particles |  | 14,729 | 10,540 | 5,262 |
| Resolution (Å)<br>(FSC = 0.143) | Overall | 3.1 | 5.8 | 8.2 |
|  | 50S subunit | - | 5.5 | - |
|  | 30S subunit | - | 6.5 | - |
| EMDB |  | EMD-33115 | EMD-33116<br>EMD-33117 | EMD-33118 |
| EMPIAR |  | EMPIAR-10985 | EMPIAR-10986 | EMPIAR-10987 |

**Supplementary Movie 1:** Slices through tomogram from dataset 3 collected on the cryoFIB-milled lamella shown in Fig. 4b (tilt series 3). The subtomogram average was mapped into the tomogram using the refined coordinates and orientations. Membrane associated and cytoplasmic ribosomes were coloured red and blue, respectively.

**Supplementary Movie 2:** Cross-correlation aligned tilt series from dataset 3 collected on the cryoFIB-milled lamella shown in Fig. 4b (tilt series 3). Scale bar: 100 nm.

